# Structural anther mimics improve reproductive success through dishonest signalling that enhances both attraction and the morphological fit of pollinators with flowers

**DOI:** 10.1101/2021.10.21.465225

**Authors:** Ethan L Newman, Katharine L Khoury, Sandy E van Niekerk, Craig I Peter

## Abstract

- Numerous studies have identified traits associated with pollen mimicry, however, the processes underlying floral deception remains poorly documented for these structures. We studied the importance of attraction and mechanical fit of anther mimics in *Tritonia laxifolia* (Iridaceae) and their relative contributions to reproductive success.
- To determine anther mimics role in pollinator attraction, we offered bees’ binary preferences to flowers painted with UV absorbent and reflecting paint. We also conducted preference experiments between flowers with excised anther mimics and unmanipulated controls, from which mechanical fit was assessed using single visits. Anther mimics effects on female reproductive success was determined using similar treatments, but on rooted plants.
- Bees preferred UV absorbent over UV reflecting anther mimics. Preference for flowers with and without the three-dimensional structures was equal. Single visits resulted in more pollen deposition on unmanipulated controls over flowers with their anther mimics excised, which was directly linked to pollen-collecting behaviour. Controls with unmanipulated anther mimics experienced more seed set than those with their anther mimics excised.
- This study provides insights into pollinator-mediated selection on deceptive floral signals and shows that three-dimensional anther mimics increases reproductive success through both attraction and pollen collecting behaviours that improves the fit between flowers and pollinators.

## Introduction

Many angiosperms exploit the perceptual biases of food-seeking visitors to obtain pollination services through traits involved in attraction, such as flower colour (Koski, 2020) and scent (Raguso, 2008). In return, pollinators receive a nutritional reward, with nectar being the most obvious. However, pollen is often overlooked as a reward but is essential for both solitary and social bees as provisions for their larvae and energy requirements (Vaudo *et al*., 2016). Pollen is also consumed by flies (Holloway, 1976; Wacht *et al*., 2000), beetles (Steiner, 1998; Steenhuisen & Johnson, 2012), birds (Coombs & Peter, 2009) and mammals, such as non-flying mammals (Melidonis & Peter, 2015; Zoeller *et al*., 2016) and bats (Herrera & Martínez Del Río, 1998; Newman *et al*., 2021). Because pollen foraging reduces male fitness, to compensate, many plants have evolved floral signalling structures that imitate pollen and deceive insects into thinking that they will receive a pollen reward, but improves the reproductive success of the plant instead (Vogel, 1975; Osche, 1979; Osche, 1983a; Osche, 1983b; Lunau, 2006). This pollen imitating signals differ from typical nectar guides in that they share yellow, UV absorbent colours between 500 to 600nm that resemble the bright yellow flavonoids and carotenoids in the pollenkitt (Harborne & Grayer, 1993). This colouration is thought to be one of the first colour signals to have evolved in the angiosperms (Lunau, 2000), which is inferred by the pollination of basal angiosperms by pollen foraging insects (Endress, 1990; Yuan *et al*., 2008; Bao *et al*., 2019; Peris *et al*., 2020).

Consequently, these pollen imitating floral signals are widely documented in the angiosperms and is found across several plant families and range from two-dimensional “guides” situated near the flower gullet to three-dimensional structures often associated with staminodes (Lunau, 2000). Pollen imitating structures have been recorded as vestigial staminodes in the Bignoniaceae (Guimarães *et al*., 2008; Milet-Pinheiro & Schlindwein, 2009), Plantaginaceae (Walker-Larsen & Harder, 2001b; Dieringer & Cabrera, 2002) and Begoniaceae (Agren & Schemske, 1991; Schemske & Agren, 1995; Schemske *et al*., 1996). Pollen mimicking labellum structures and colours have been documented in a range of Batesian mimicry systems in the Orchidaceae, pointing to the importance of such structures in successfully deceiving pollinators (Nilsson, 1983; Peter & Johnson, 2008). Additional evidence inferred from a recent case study shows that pollen imitating structures occurs in 28% of all angiosperm flora of the Alps (Lunau *et al*., 2017), which suggests that its occurrence may be under-documented.

Despite their prevalence, the selective advantage of pollen imitating structures remains understudied. Studies investigating the occurrence of pollen mimicry, primarily document their presence or spectral reflectance properties and few investigate the underlying evolutionary processes. To date, studies have focussed on the functional role of vestigial staminodes (Walker-Larsen & Harder, 2001b), including their role in attracting pollinators and improving the morphological fit of pollinators when visiting flowers (Walker-Larsen & Harder, 2001b; Dieringer & Cabrera, 2002; Guimarães *et al*., 2008; Milet-Pinheiro & Schlindwein, 2009). These studies compare components of fitness between flowers with their staminodes excised against unmanipulated controls, often considering handling time as an additional fitness surrogate. However, it remains unclear whether improved fitness on unmanipulated controls is the result of the three-dimensional structure of the pollen mimic creating a hindrance to pollinators accessing the nectar reward (Martos *et al*., 2015), or whether reproductive success is improved when pollen-collecting behaviour is focussed on the anther mimics rather than the functional anthers.

Demonstrating the inability of the pollinator (signal receiver) to discriminate between the signals of the model (pollen) and the mimic (pollen mimic) is a crucial line of evidence for inferring floral mimicry (Newman *et al*., 2012; Schiestl & Johnson, 2013), including pollen mimicry (Lunau, 2000). Although preferences of pollen imitating structures have been documented (e.g., Duffy & Johnson, 2015; Milet-Pinheiro & Schlindwein, 2009; Lunau, 2014), the lack of evidence for pollen-collecting behaviour by bees on pollen imitating structures remain surprising. This is because pollen collection is an essential behaviour of both social and solitary bees, both of which require pollen as nutritional resources for their brood, and such pollen imitating structures are likely adaptions to exploit such behaviour. Nevertheless, Vogel (1978) observed differences in the way that bees handle anther mimics over stamens. His observations indicate that bees search for pollen using specific behaviours that they do not exhibit when interacting with anther mimics. However, it is not determined whether it is experienced bees that exhibit these behaviours or not, leaving the question open on whether bees exhibit “true” pollen-collecting behaviour on anther mimics (*see* Lunau *et al*., 2017 for details). It is also unknown whether pollen-collecting behaviour occurs on both three-dimensional anther mimics and two-dimensional pollen imitating colour markings. For example, Ellis and Johnson (2010) found that three-dimensional floral signals in *Gorteria diffusa* elicited a higher proportion of mating behaviours by male bee flies, *Megapalpus capensis*, resulting in higher pollen export in deceptive morphs with three-dimensional floral signals, compared to varieties with two-dimensional floral signals. This behaviour is thought to be elicited by a tactile association between the three-dimensional structure and the pollinator. Indeed, similar outcomes are likely to occur in flowers with pollen imitating structures; with increased pollen collecting behaviour on three-dimensional pollen mimics versus two-dimensional pollen imitating markings.

*Tritonia* (Iridaceae) is a small African endemic genus of 30 species with pollen imitating structures ranging from those species having no anther mimics and “unicolour” tepals to two-dimensional pollen imitating colour markings contrasting against the tepal colours, and three-dimensional anther mimics formed by raised structures projecting from the tepals, contrasting strongly with the tepals (**Fig**. S1). *Tritonia laxifolia* Bentham ex Baker (Iridaceae) (Fig. **1a**) is one such species with prominent three-dimensional anther mimics on its lower lateral and median tepals that appear as yellow teeth-like structures (Fig. **1b**). Using colour vision analysis combined with binary preferences, single visits, and video recordings, we explore both sensory and morphological fit of anther mimics with pollinators in this system, including their relative contribution to fitness. We ask the following broad questions. 1. Do pollinators prefer the colour and structure of anther mimics? 2. Do three-dimensional anther mimics facilitate the morphological fit of pollinators with flowers? 3. Do preferences and morphological fit of pollinators to flowers with structural anther mimics have consequences for female reproductive success?

**Figure 1.**
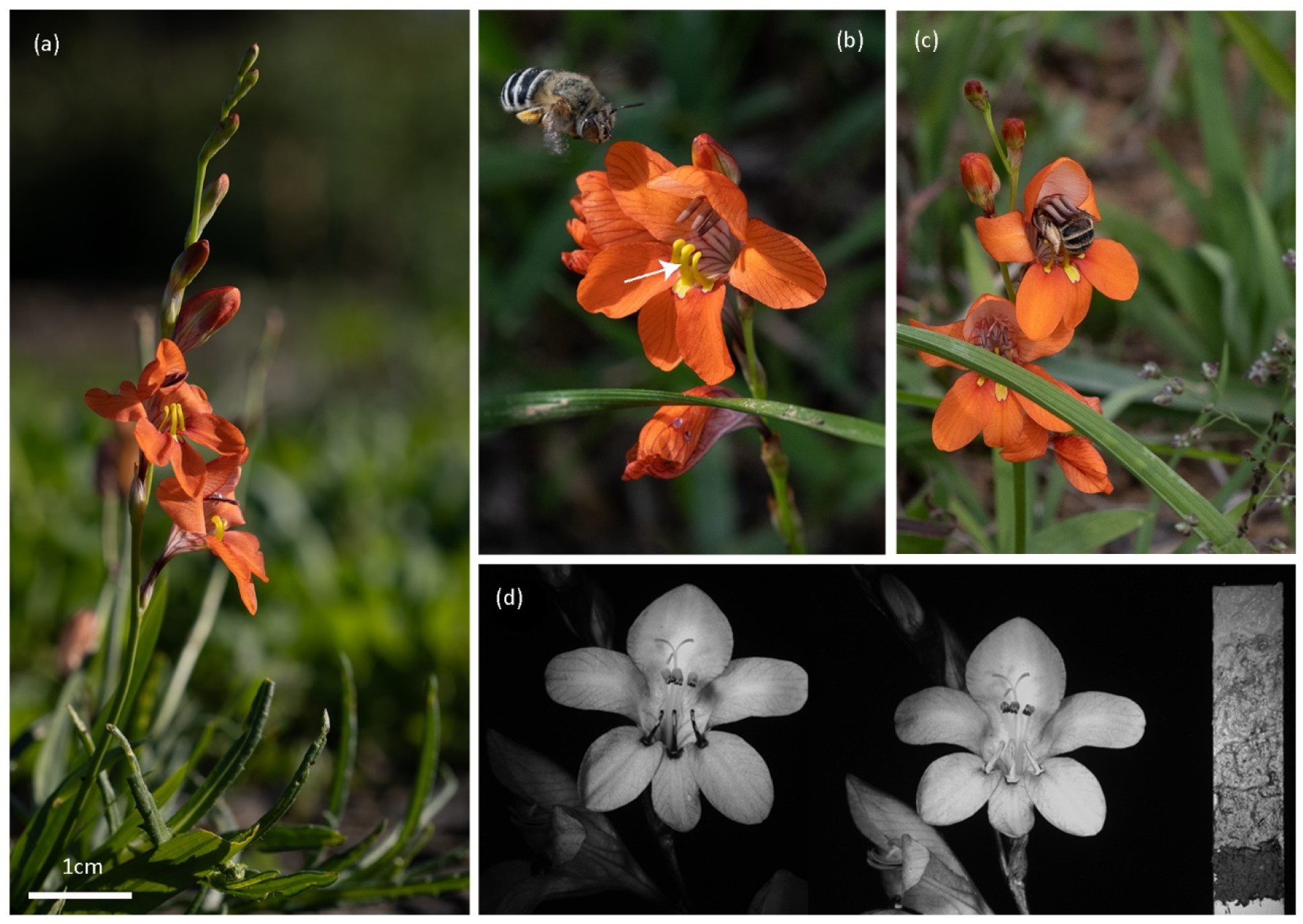
Colour plate of the study system. (a) *Tritonia laxifolia* (Iridaceae) in flower at Kwa-Pikoli, Fish River Pass. (b) *Amegilla fallax* approaches a flower, the white arrow highlights three-dimensional anther mimics on each lower tepal. (c) Bee visitor required to crawl onto and over the anther mimics to contact reproductive parts of the flower. (d) UV images of an unpainted control (left) and a flower with its anther mimics painted with UV reflecting orange paint.

Specifically, in question 1, we determine whether pollinators prefer yellow painted UV absorbent anther mimics over orange painted UV reflecting anther mimics, and whether pollinators prefer three-dimensional anther mimics over two-dimensional pollen imitating markings. Furthermore, we expect that anther mimics should contrast strongly with all floral traits and that the anthers should be camouflaged against the dorsal tepals and be indistinguishable to bees. For question 2, we determine if bees transfer more pollen onto virgin stigmas of flowers with their anther mimics unmanipulated compared to when they are removed and determine if this is the result of pollen collecting behaviours. For question 3, we determine whether unmanipulated flowers with anther mimics present set more seed than flowers with anther mimics removed in a selection experiment that uses the same experimental procedures as questions 1 and 2.

## Materials and Methods

### Study species and localities

*Tritonia laxifolia* (Iridaceae) Bentham ex Baker is a small, deciduous, winter growing geophyte which flowers from March to June in disturbed habitats along the east coast of Africa from Port Elizabeth in the Eastern Cape of South Africa to Tanzania (de Vos, 1982). The scentless, zygomorphic flowers are orange red with the adaxial surface of the dorsal tepal being a contrasting pale pink. The most striking feature of the flowers are the three peculiar bright yellow three-dimensional structures on each of the lower tepals referred to here as anther mimics (Fig. **1a**). In addition, *T. laxifolia* has three inconspicuous, light pink coloured anthers. Receptive stigmas are deeply divided with three style branches becoming recurved and coarsely pustulate when receptive (Manning *et al*., 2002). Flowers typically last between two and three days and are protandrous with distinct male and female phases (Ethan Newman personal observation).

Our study was conducted from April to June in 2019 and 2021, near Fish River Pass, Kwa-Pikoli (−33.241132°, 27.014889°), (**Fig**. S2a) and Mosslands farm 18 km south west of Makhanda/Grahamstown (−33.401357°, 26.432470°), Eastern Cape, South Africa. Here, *T. laxifolia* occurs on seasonally wet clay in disturbed thicket vegetation dominated *Euphorbia tetragona, E. triangularis* and *Aloe ferox*. At both study sites, *T. laxifolia* is primarily visited by *Amegilla fallax* (Fig. **1b, c**), with *Apis mellifera scutellata* and pollen-collecting bees present in lower abundance. Medium and large butterflies, *Colotis eris eris, Pinacopteryx eriphia eriphia* and *Papilio demodocus* were abundant and frequently visited the flowers (**Fig**. S2b, c).

#### 1. Do pollinators prefer the colour and structure of anther mimics?

##### Colour preferences

To determine if the yellow UV absorbent colour of anther mimics is important in attraction, we conducted binary preferences at Mosslands between the 7th and 23^rd^ of May 2021, between 0900hrs and 1400hrs at temperatures consistently exceeding 20°C. Fresh flowers were picked from the field before pollinators arrived. We removed the yellow UV absorbent colour signal from one-half of the flowers by painting the anther mimics with orange UV reflecting paint (Dala Neon Orange) that is similar in colouration to the adjacent tepals. We mimicked the yellow UV absorbent colour of the anther mimics by painting the anther mimics of the other half of the experimental flowers with UV absorbent yellow paint (Dala craft paint, yellow). Paints were applied to the entire UV absorbent yellow part of the pollen mimic using a #7 insect pin at least an hour before preference experiments started and allowed to dry. To control for the potential influence of the scent of the paints, an equal amount of paint from the opposite treatment was applied to the inner part of the container serving as a vase to hold the inflorescences. Binary colour preferences included two experimental trials: 1. Anther mimics painted with yellow UV absorbent paint versus orange UV reflecting paint. 2. Anther mimics painted with yellow UV absorbent paint versus unpainted anther mimics. Experiment 1 tests the importance of the UV absorbent yellow in attracting the pollinator. Experiment 2 is a control that assesses whether the pollinators are equally attracted to the yellow UV absorbent paint applied in experiment 1 and unpainted anther mimics.

We used the bee interview technique for both experiments [e.g. Johnson *et al*. (2003)], as the flowers were too numerous within the population to wait for bees to approach stationary arrays. In these experiments, flower pairs were suspended in two 25ml tubes filled with water and fixed at the end of a bamboo stick (±2 m), arranged approximately 10 cm apart. Preferences were executed by placing pairs near a foraging pollinator, and pollinators were offered a binary preference. We recorded the insect species and the individual pollinators first preference. Statistical differences within experimental treatments were analysed using generalised linear mixed-effects models (GLMMs) with binomial error distributions and logit link functions, where treatments were assigned as fixed factors and binary preferences as the response. Because it was challenging to swap inflorescences within a pair after each visit to account for non-independent positioning of pairs, specific pairs (i.e., different preference sticks containing choices) were incorporated as a random factor.

##### Preference for the physical structure of the anther mimics

We picked a total of 128 inflorescences in bud or the early stages of flowering over 17 days between 20th of May and the 06th of June 2019. As inflorescences matured, we emasculated flowers using a pair of fine forceps to prevent pollinator preferences from being influenced by flowers in different stages of anthesis (e.g., pollinators may prefer male phase flowers containing pollen over female phase flowers). To determine the effects of anther mimics on pollinator preference, we carefully excised all three anther mimics from all available flowers from exactly half of the experimental inflorescences (n=64) using a surgical blade. The other half remained unmanipulated (only the anthers were removed) (n=64). We refer to these inflorescences/flowers as “anther mimics excised” and “unmanipulated controls” throughout the manuscript (**Fig**. S2d). Essentially, the excision of the anther mimic removes the physical structure, but the round yellow mark on the tepal that remains after excision serves as a two-dimensional visual component of the anther mimic. These inflorescences were kept in a cool room at 10°C until the stigmas became receptive. Once receptive, experimental inflorescences were used to disentangle the function of the three-dimensional structure of the anther mimic, using experiments that simultaneously test the roles of visual signalling and morphological fit (pollen deposition) in the pollination process, as explained below.

To test whether the physical structure of anther mimics is associated with pollinator attraction, we conducted preference experiments over four days between the 29^th^ of May and 04^th^ of June 2019 between 09:00 and 14:00 depending on pollinator activity. Inflorescences with anther mimics excised and unmanipulated controls were organised into ten pairs with individuals placed approximately 10 cm apart and spaced about 25 cm from other pairs. These pairs were arranged at the same height relative to naturally occurring flowering plants within the population, and the control and experimental inflorescences were matched for the number of open flowers (either 1 or 2). Once a visitor entered the arena, one of the authors recorded the binary preference made by visitors to treatments within a pair. Observers also recorded the sequence of visits to pairs by each pollinator individual. Only first choices were included in the statistical analysis (switches to alternative phenotypes within a pair were excluded from the preference analysis and only used in the single visit experiments). All pollinators were incorporated in the data analysis. We treated binary preferences as the response and treatments as fixed factors in a GLMM that considered a binomial error distribution and a logit link function, with pollinator individual treated as a random factor to account for non-independence in the data resulting from individual behaviour.

To determine whether bees can perceive differences between the anther mimics, anthers, pollen and adjacent flower tissue including paints used in preference experiments, we measured colours from different segments of the flower involved in attraction from between four to 13 receptive female phase flowers from different individuals from Makhanda. We separated anther mimics from the anthers, pollen, dorsal and lateral sepals using a surgical blade. We also painted the central pollen mimic of ten individuals with UV reflecting orange paint and four individuals with UV absorbent yellow paint used in colour preferences. Once dry, these were measured, together with each flower segment over the UV-visible range between 300 to 700nm using an Ocean Optics S2000+ spectrometer with a DT-mini light source and fibre optic probe (UV/VIS 400μm). To assess the qualitative pattern of UV absorbing and UV reflecting parts of the flower, we photographed flowers using a UV camera (*Methods S1*).

Spectra was then imported into bee colour space (Chittka *et al*., 1992) using hyperbolically transformed quantum catches and reflectance spectra of foliage occurring within the immediate vicinity where preference and selection experiments were conducted (*see* **Fig**. S2a for an image of the habitat). We did not use the main attractive surface of the flower as background colour (either lateral tepal or dorsal tepal), as bees make consistent decisions for experimental flowers separated by 10cm (see results) which means that actual preferences are considered with vegetation as the background. Standard D65 daylight illumination was used. To determine whether 1) bees could perceive differences between the anther mimics and adjacent floral structures, and 2) between the anthers and adaxial surface of the dorsal tepal, mean Euclidean distances with bootstrapped 95% confidence intervals were determined as chromatic contrasts. This was obtained using the *bootcoldist* function implemented in the R package “*pavo*” (Maia *et al*., 2019). Colour distances below the perceptual threshold of 0.11 hexagon units is considered as indistinguishable by pollinators (Dyer, 2006; Bukovac *et al*., 2017).

#### 2. Do anther mimics facilitate morphological fit of pollinators with flowers?

##### Physical structure of the anther mimic on pollen deposition

Single visits to virgin flower were used to test whether anther mimics enhance pollen deposition to receptive stigmas. We were able to directly link pollinator preferences for the physical structure of anther mimics (see the previous section) with pollen deposition to treatments in the following manner: After each foraging bout, the second observer identified the “preferred” flowers in the experimental array that received a single visit and carefully removed the stigmas of visited flowers near the base of the ovary. Stigmas were immediately placed in a labelled 2.5 ml centrifuge tube and maintained in an ice filled cooler box while in the field. Experimental arrays were immediately reconstructed with fresh inflorescences maintaining a constant ten inflorescence pairs all with virgin flowers. Once a new inflorescence was introduced, the positions of the treatment was swapped. Stigmas were embedded in heated fuchsin gel mounted on microscope slides later the same day. A dissecting microscope was used to count the total number of *T. laxifolia* pollen grains on each stigma. *T. laxifolia* pollen was easily identified relative to other community members represented in a pollen library of the site.

To account for the high number of zeroes in the dataset, that led to overdispersion in an initial model using a Poisson error distribution, we used a GLMM with a negative binomial error distribution and log link function (Zuur *et al*., 2009; Brooks *et al*., 2019). In our analysis, we removed butterflies from the full dataset, only retaining bees as the primary pollinators, although we report on both [i.e., butterflies act as nectar thieves within the Fish River Pass population]. Treatments: flowers with their anther mimics removed and unmanipulated controls species were assigned as fixed factors, and *T. laxifolia* pollen counts were treated as the response. Individual visitors were treated as random factors to account for non-independence resulting from similar pollinator morphology and behaviours.

##### Physical structure of the pollen mimic on pollen-collecting behaviour

To assess whether pollen deposition from single visits is the consequence of pollen-collecting behaviour exhibited by bees on anther mimics. We extracted behavioural data from 159 videos recorded during our field season at Mosslands. We set up arrays similar to the experiments investigating morphological fit described above. This yielded 43 videos of bees on flowers with the anther mimics excised and 116 on flowers as unmanipulated controls. These videos were recorded using a Canon 5D MKIV with a 100 mm USM macro lens shot at 30 fps.

GLMMs with a binomial error distribution and a logit link function was calculated to determine a statistical difference in pollen-collecting behaviour. Three separate models were run, namely, the presence or absence of “scraping” and “pulling” on anther mimics among treatments, including the “proportion pollen-collecting behaviour that resulted in contact with the reproductive parts” between treatments. In these models, bee individual was included as a random factor to account for non-independence regarding individual behaviour.

#### 3. Do preferences for, and morphological fit on anther mimics have consequences for seed set?

To determine whether anther mimics are associated with female reproductive success. We compared seed set from treatments with their anther mimics excised from unmanipulated controls of naturally occurring rooted plants within the population. Similar to the previous two experiments, we emasculated all experimental flowers and excised anther mimics from a total of 56 flowers (anther mimics removed) and left anther mimics of 51 flowers intact (unmanipulated controls). Providing a total of 107 experimental flowers. We covered 28 individuals with their anther mimics removed and 34 unmanipulated controls (62 treatments) with 33 chicken wire boxes rooted with steel tent pegs [from a total of 51 inflorescences]. We did this to determine if butterflies made a significant contribution to fitness. If this is the case, we expect open treatments to experience a higher proportion seed set than caged individuals. Wire cages had holes large enough to allow bees [*A. fallax* body length: distance from head to tip of abdomen, 11.07±0.97mm (4)] to enter the cages (25mm holes), (*see* Fig. S2e), but small enough to prevent white butterflies from entering (EN and SVN personal observation). The remaining 32 inflorescences were left uncaged, containing 28 manipulated and 17 unmanipulated flowers (45 treatments). After three weeks, we collected fruits and discerned fertilised from aborted ovules. Fertilised ovules were much larger, hard, and green in appearance whereas aborted ovules were smaller, soft, and shrivelled in appearance.

Of the 33 cages setup initially, five were destroyed by cattle, leaving a total of 28 cages intact. Fruit set from the five destroyed cages were discarded from the analysis. Statistical differences in the proportion seed set amongst treatments (anther mimics removed and unmanipulated controls) and exposure (caged versus open) and their interaction were calculated using GLMM with a beta-binomial error distribution and a logit link function. Individual inflorescences were treated as a random factor to control for multiple treatments per inflorescence.

All GLMMs were calculated using the *glmmTMB* command from the package “*glmmTMB*”, significance of fixed effects were determined using the ANOVA, type III command from the “*car*” package (Bolker *et al*., 2009) and contrasts among interaction terms for the selection experiment was determined using the *emmeans* command from the package “*emmeans*”. All models were checked using the “*DHARMa”* package. In the process, we discovered that the final model was overdispersed, and we corrected for overdispersion using a beta-binomial error distribution to model the proportion seed set from the selection experiment (Harrison, 2015). Median values and 90% confidence intervals for plotting were obtained from model predictions using 1000 bootstrap samples calculated using the *bootMer* command from the package “*lme4*”. All data analysis was conducted using the R statistical environment (R Development Core Team, 2021).

## Results

### 1. Do pollinators prefer the colour and structure of anther mimics?

#### Colour preferences

At Mosslands, 14 *A. fallax* bees showed a significant selection bias for flowers with anther mimics painted with yellow UV absorbent paint over flowers painted with orange UV reflecting paint (χ^2^=11.00, df=1, *P* <0.001, Fig. **2a**). The loci of UV reflecting orange paint being close to the loci of the orange of adjacent flower tepals in bee colour space (Fig. **1d**, **3;** Fig **S3**, **S4a**). In contrast, 13 *A. fallax* bees made equal choices between anther mimics painted with yellow UV absorbent paint over unpainted controls (χ^2^=0.15; df=1, *P* =0.70, Fig. **2b**), the loci of both these yellow colours clustering together in bee colour space (Fig. **3**, **S4b**). Chromatic contrasts reveal that anther mimics were above the threshold of discrimination of 0.11 hexagon units (Dyer, 2006; Bukovac *et al*., 2017) when compared to all other floral traits (Fig. **3**. **S3**, **S4c**). However, the anthers were perceptually similar to the adaxial surface of the dorsal tepal, being well below the threshold of 0.11 hexagon units (Fig. **S4d**).

**Figure 2.**
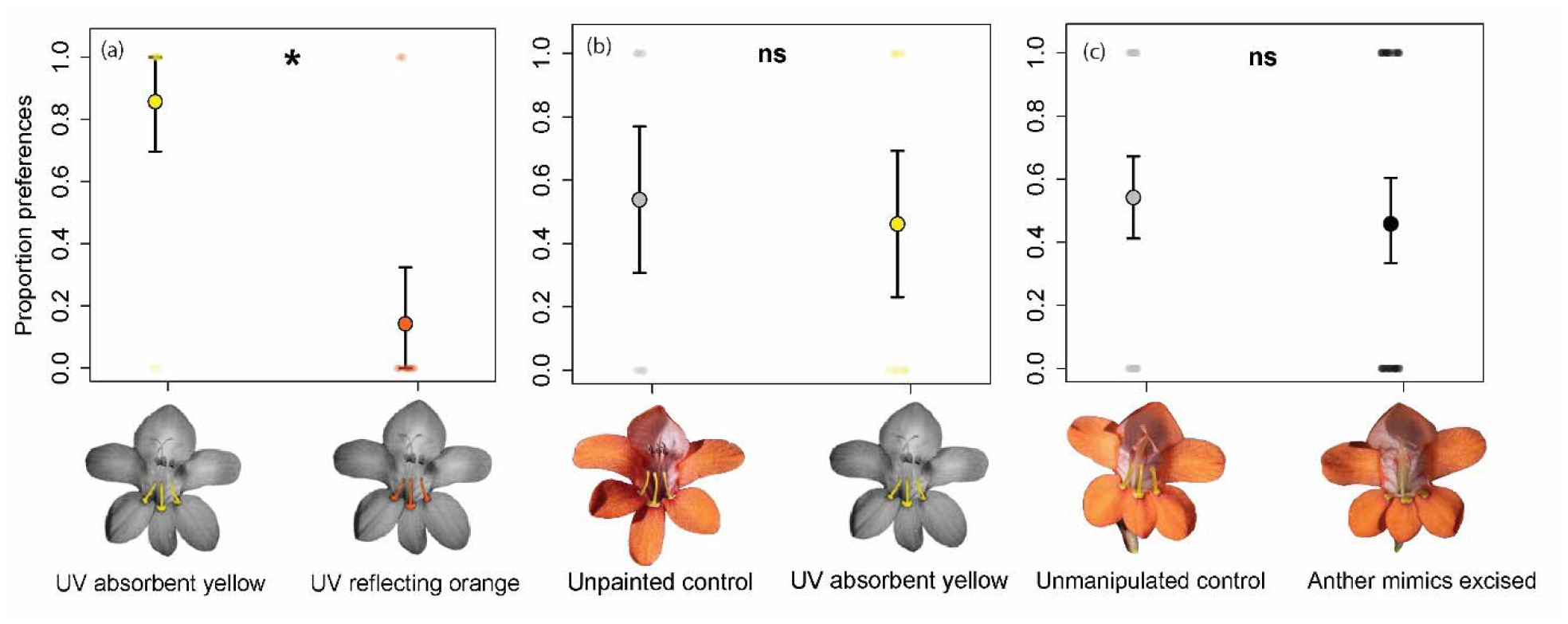
Binary preferences of bees based on their first choices (a) between anther mimics painted with yellow UV absorbent paint and orange UV reflecting paint; (b) choices between unpainted controls with naturally yellow anther mimics and anther mimics painted with yellow UV absorbent paint; and (c) choices between unmanipulated controls and flowers with anther mimics excised. Coloured circles represent median model predictions, error bars refer to 90% confidence intervals for model predictions and small coloured points represent balanced binary preferences for each treatment. *ns=P*>0.05, *=*P*<0.05.

**Figure 3.**
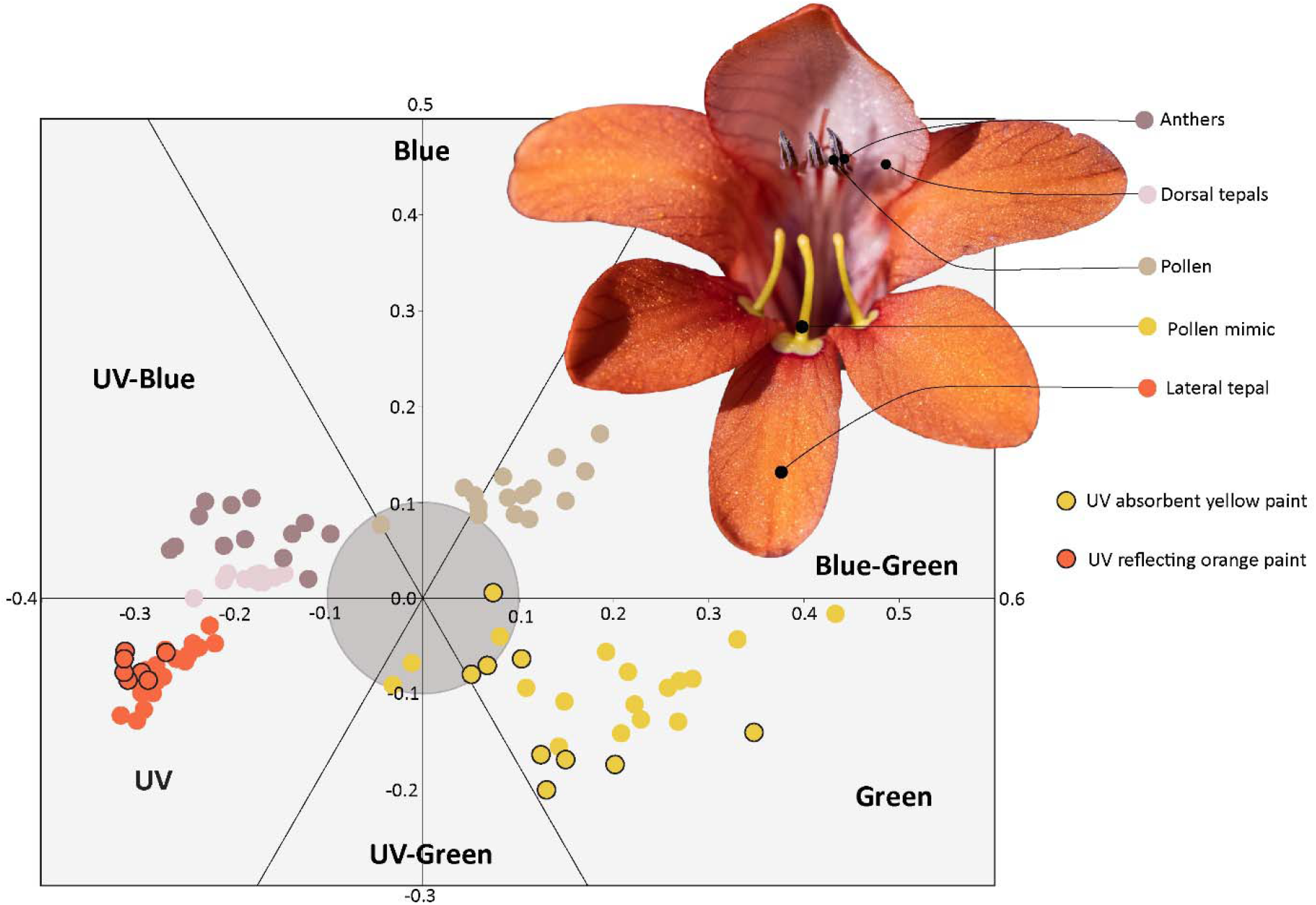
Colour spectra from different parts of the flower plotted in bee colour space. Colours represent actual colours from respective parts of the flower as perceived by humans. Points with a black outline are measured from anther mimics painted with either orange UV reflecting paint or yellow UV absorbing paint. Spectra in the central grey circle appear achromatic to bees (0.1 hexagon units).

#### Preference for the physical structure of the anther mimic

At Fish River Pass, we interviewed 40 insects, of which 20 were bees (16 *A. fallax* and four *A. mellifera scutellata*), and 20 were butterflies (19 *C. eris eris* and one *P. eriphia eriphia*), which made a total of 88 first preferences. The model including all insects (both bees and butterflies) χ^2^ =3.81, df=1, *P* =0.051 showed no significant preference for flowers with or without physical anther mimics present. Removing butterflies from the dataset, did not alter this result and bees alone showed no preference for flowers with or without anther mimics (χ^2^=0.66, df=1, *P* =0.415, Fig. **2c**).

### 2. Do anther mimics facilitate morphological fit of pollinators with flowers?

#### Physical structure of the anther mimic on pollen deposition

At Fish River Pass, we obtained a total of 74 single visits from 34 insects. Of these, 41 visits were made by bees (14 *A. fallax* and two *A. mellifera scuttelata*) and 33 by butterflies (17 *C. eris eris* and one *P. eriphia eriphia*). Pollinators deposited significantly more pollen grains onto virgin stigmas of unmanipulated controls, compared to flowers with their anther mimics removed (χ^2^ =6.70, df=1, *P* =0.009). This result did not change when butterflies were removed from the dataset, which highlights the significant contribution of bees to pollen deposition in the experiment (χ^2^=5.74, df=1, *P* =0.017, Fig. **4**).

**Figure 4.**
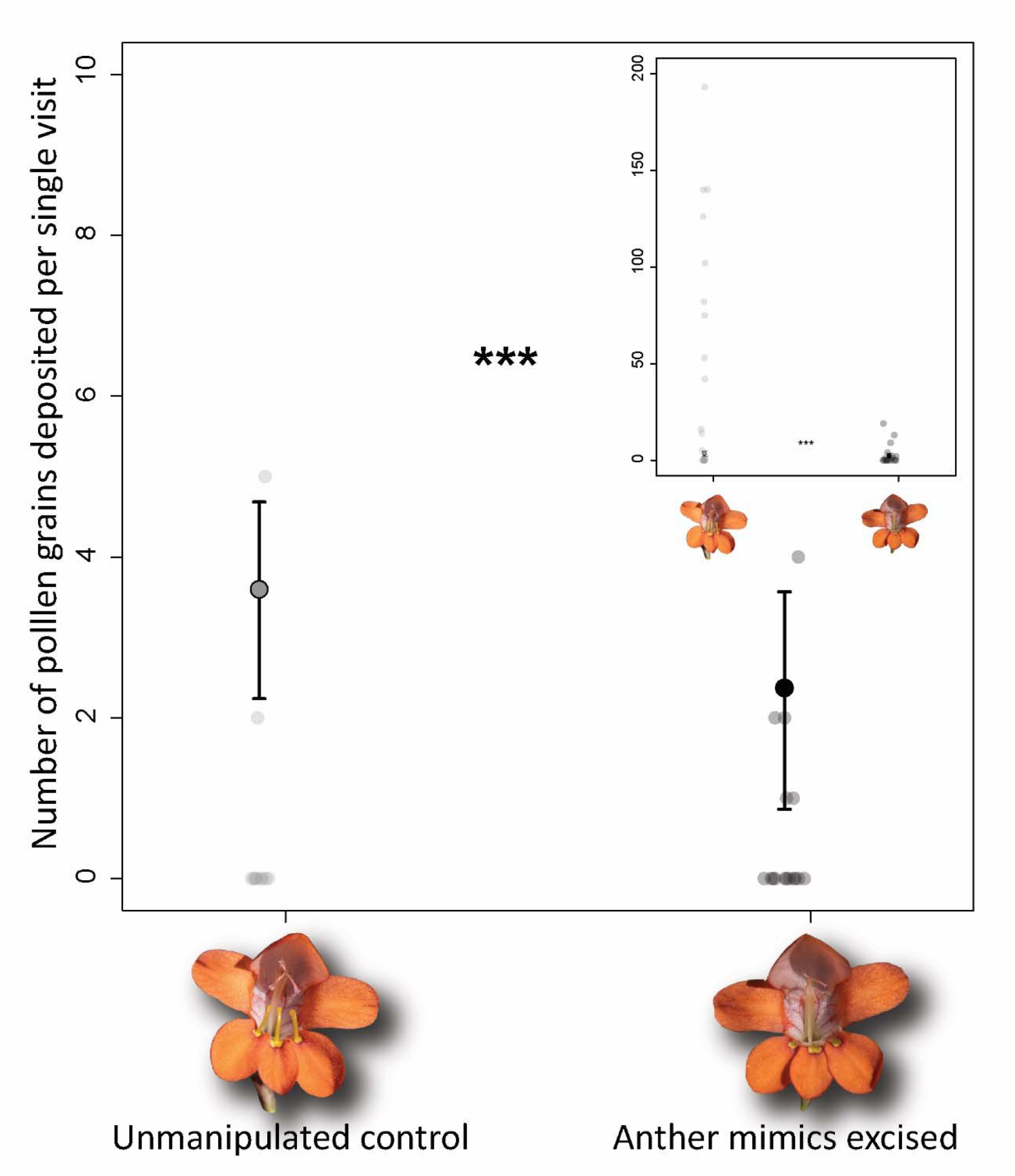
Unmanipulated flowers of *Tritonia laxifolia* with anther mimics present, received significantly more pollen deposited on their stigmas following a single visit by a pollinating bee compared to flowers with their anther mimics excised. Coloured circles represent median model predictions, error bars refer to 90% confidence intervals for model predictions and small points represent the number of pollen grains deposited for each single visit replicate. Inset shows full extent of data points. *** *P*<0.001.

#### Importance of anther mimic structure for pollen-collecting behaviour

At Mosslands, we recorded 155 *A. fallax*, a single Allodape and a single Halictid bee visiting 42 flowers with the anther mimics excised and 115 unmanipulated control flowers. Bees demonstrated a higher proportion of pollen-collecting behaviour on anther mimics (57.14%),(χ^2^=10.14; df=1, *P* =0.001). They exhibited more scraping and pulling behaviour on anther mimics of unmanipulated controls, compared to when they were excised (scraping: χ^2^=8.85; df=1, *P* =0.002, Fig. **5a**; pulling: χ^2^=10.10, df=1, *P* =0.001, Fig. **5b**).

**Figure 5.**
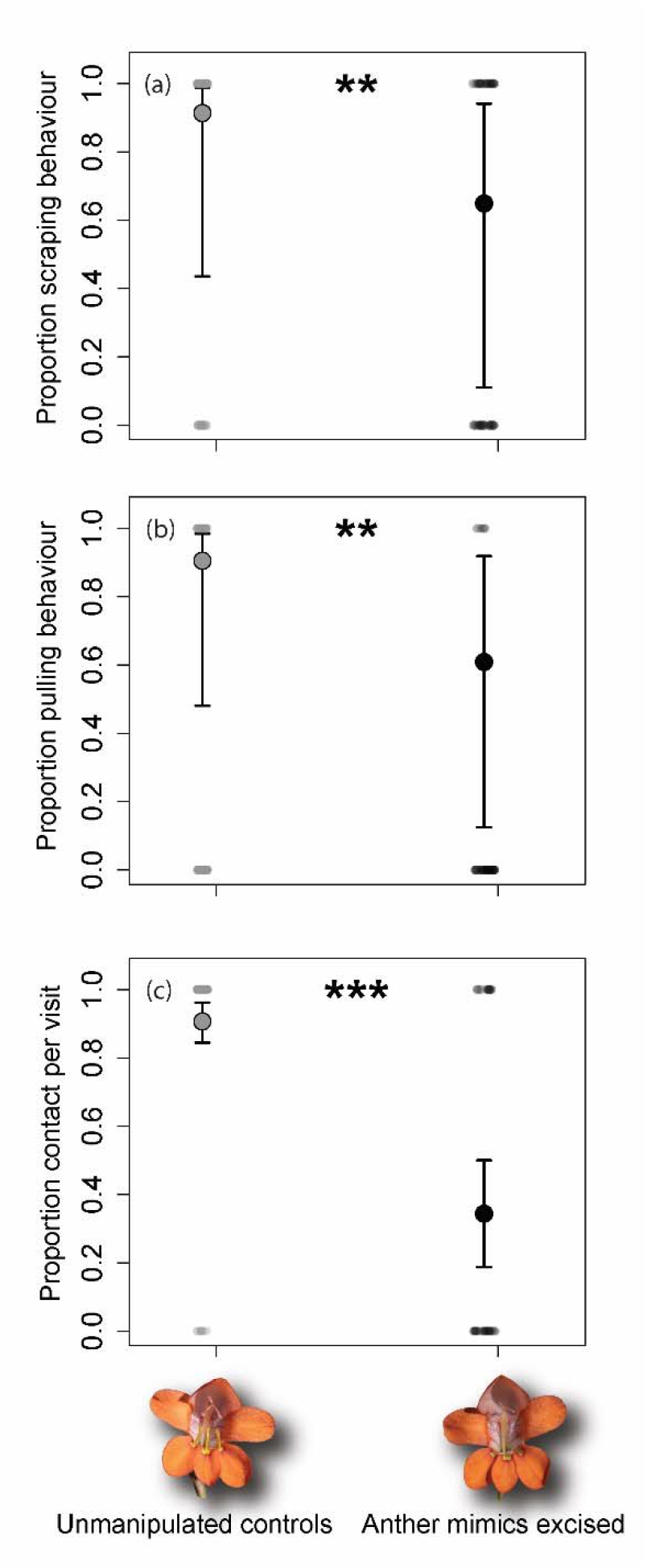
Pollinator behaviour recorded on unmanipulated controls compared to flowers with anther mimics excised. (a) Proportion of visits with bee scraping anther mimics, (b) Proportion of visits with bees pulling at anther mimics, (c) Proportion of visits by pollen collecting bees (either scraping or pulling behaviour) contacting reproductive organs. Coloured circles represent median model predictions, error bars refer to 90% confidence intervals for model predictions and small points represent binary outcomes. (i.e., presence or absence of behaviour exhibited). **=*P*<0.01*** *P*<0.001.

#### Contact with reproductive parts and associated pollen-collecting behaviour

The proportion of bees making contact with anthers and stigmas was significantly higher on unmanipulated controls versus flowers with anther mimics excised (χ^2^=30.87, df=1, *P* <0.001, Fig. **5c**). The result remained similar when nectar foraging bees were excluded and only pollen collecting bees considered (χ^2^=11.62, df=1, *P* <0.001).

### 3. Do preferences for, and morphological fit on anther mimics have consequences for seed set?

Butterflies did not contribute to seed set at the Fish River Pass site, seed set of the plants caged to exclude butterflies being similar to controls (χ^2^=0.33, df=1, *P* =0.69, Fig. **6**). However, the removal of the physical anther mimics led to a significant decrease in seed set (χ^2^=10.90, df=1, *P* <0.001, Fig. **6**). There was no significant interaction between pollinator exclusion and pollen mimic excision, and none of the contrasts between caged and open treatments was significant. (χ^2^=0.08, df=1, *P*=0.78, Fig. **6**).

**Figure 6.**
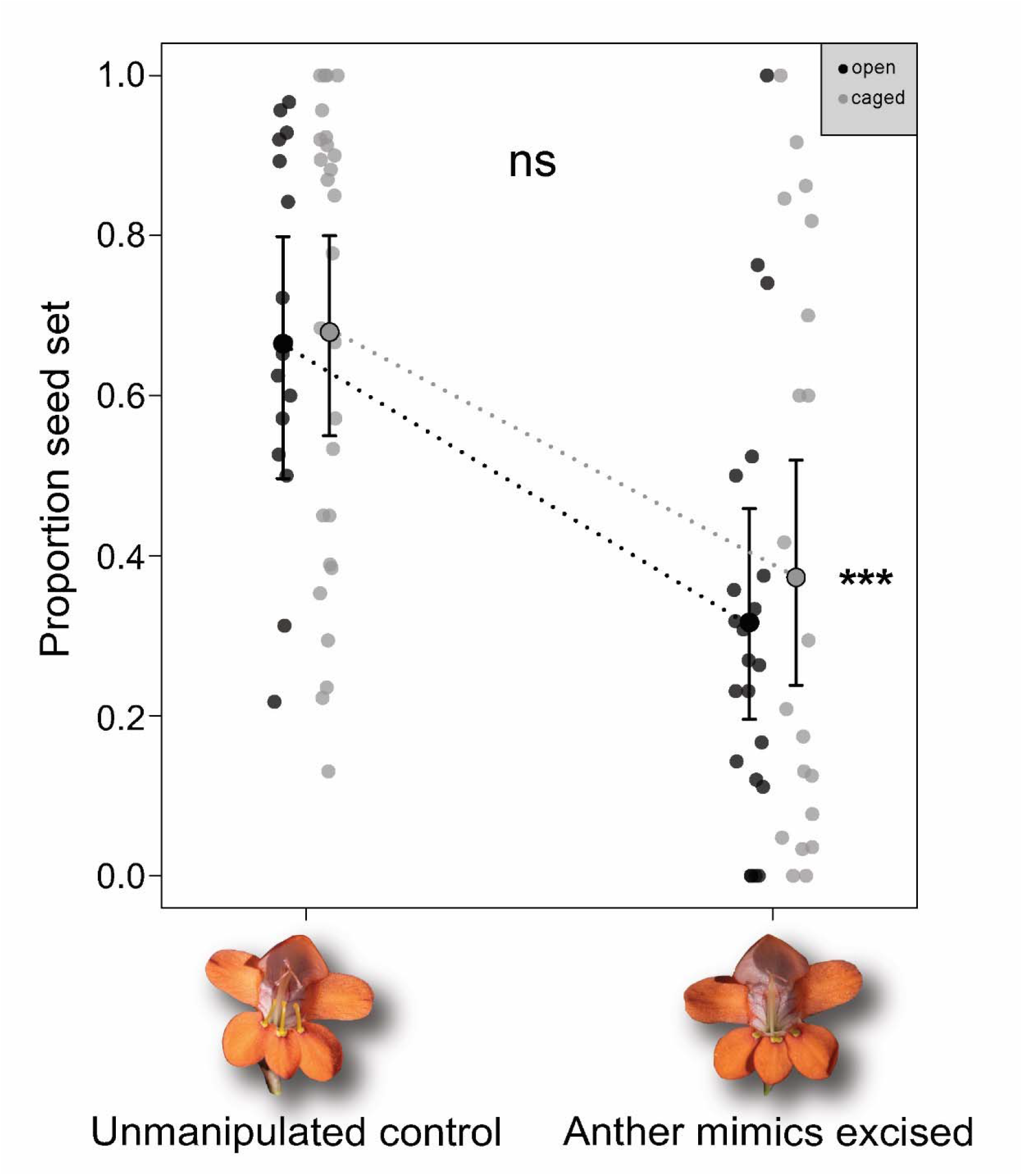
Mean proportion seed set from caged and open treatments on rooted unmanipulated controls and flowers with their anther mimics removed. Manipulated caged and open treatments experience significantly lower seed set compared to unmanipulated caged and open treatments. However, there is no significant difference in seed set within manipulated and unmanipulated treatments for caged versus open treatments, suggesting that the presence of butterflies in open treatments did not contribute significantly to seed set. ** *P*=0.001.

## Discussion

*Tritonia laxifolia* accomplishes pollination through floral mimicry that deceives the sensory abilities of the primary bee pollinators through both crypsis, as well as generalised pollen mimicry. The functional anthers are concealed against the dorsal sepal while physical, three-dimensional anther mimics deceive pollen collecting bees, focusing their attention on the lower tepals at the entrance to the flower. Besides the importance of both these visual and tactile signals, our data show that the three-dimensional anther mimics play a crucial role in precisely positioning pollinators to deposit pollen on stigmas and presumably remove pollen from anthers

Yellow UV absorbing floral signals are considered important in orientating pollen foraging insects to flowers (Lunau, 2014), and laboratory experiments using untrained naïve bumblebees (*Bombus terrestris*) demonstrate preferences for the visual signals of anther mimics by orienting themselves towards the pollen signal of dummy pollen and touching the mimics with their antennae [*see* Lunau (2000), Fig. 4 and 10]. Bees in our study selected flowers with anther mimics painted with yellow UV absorbent paint, exclusively over flowers with mimics painted with orange paint that reflected UV. The paints used for these manipulations approximate the respective floral parts in bee colour space, and the bees are unlikely to be able to distinguish the UV absorbent yellow paint from the unpainted yellow anther mimics, or the orange UV reflecting paint from the tepals in preference experiments. This is supported by mean Euclidean distances with confidence intervals that are either less than, or overlaps with the perceptual threshold of 0.11 hexagon units (Dyer, 2006; Bukovac *et al*., 2017). Similar experiments altering the UV colour signal by applying sunscreen to flowers resulted in reduced preferences by bees (Johnson & Andersson, 2002), which may have consequences for reproductive success. However, we suspect that the extreme effects of UV alteration in our study that led to absolute preferences to unpainted controls, may have been the result of bees specifically foraging for pollen rewards as indicated by a higher proportion of pollen collecting behaviour on unmanipulated controls compared to nectar foraging (*see* **Fig**. S5).

In bee colour space, the colour of the pollen and anthers contrasted strongly with that of all other floral parts except the pink dorsal tepal directly behind the anthers (**Fig**. S4). We interpret this as a case of crypsis to prevent pollen collecting insects from discovering the anther and reducing male fitness by collecting pollen as a reward (*see* Xiong *et al*., 2019). By camouflaging the pollen against the background of the dorsal tepal, *T. laxifolia* deceptively directs attention to the yellow UV absorbing signal of the anther mimics, making bees likely to ignore the actual pollen of the flower, at least in inexperienced individuals. We have however observed that bees do collect pollen from flowers by pushing their head against the anthers and grooming themselves directly afterwards, often occurring following an attempt to remove pollen from the anther mimics (*Video S1*). Together with the nectar reward, this behaviour may act as a trade-off to ensure that bees return to flowers, and it is likely that this occurs later in the season by more experienced bees. However, more research is required to confirm this notion.

In contrast to our findings, Duffy and Johnson (2015) showed that yellow anther mimics and pollen are virtually indistinguishable to bees. In their system, this convergence of colour may have evolved to increase the display of the pollen reward and increase visitation time on flowers which may improve reproductive success. This idea is supported in the removal of anther mimics resulting in decreased preferences to flowers with excised anther mimics and consequently seed set. When we provided pollinators choices between flowers with the anther mimics removed and unmanipulated flowers, pollinators did not show any preference. This was because the excision of the pollen mimic did not remove the yellow UV-absorbing pollen signal but made it two-dimensional instead. Therefore, we found no significant preference for the three-dimensional structure of the pollen mimic. Similar experiments have been conducted on *Jacaranda rugosa* (Bignoniaceae) by Milet-Pinheiro and Schlindwein (2009) that show decreased visitation to flowers with their staminodes excised. However, in their study, the excision of the staminode removes the yellow UV absorbent signal which is comparable to our first experiment where we painted the anther mimics the same colour as the tepals which resulted in no visits by pollinators.

Despite the lack of choices made to the three-dimensional structure of the anther mimic, pollinators transferred significantly more pollen per single visit on the stigmas of unmanipulated controls versus flowers with anther mimics excised. This is, in part, the result of the three-dimensional structure of the anther mimic that decreases the width of the flower entrance between the anther mimic and the reproductive part of the flower. Preliminary observations by the authority on the genus state that “The function of the calli (anther mimics) is probably to diminish the space in the throat of the perianth, thus ensuring that a visiting insect will brush with its back against the anthers and stigmas” (de Vos, 1983). Indeed, the distance between the closest stigma branch and the top of the anther mimic on the median tepal is 3.45±0.19mm (n=20), which is 1.28mm less than the thorax height of the most abundant bee pollinator *A. fallax* 4.73±0.07mm (n=12) (*Methods S2*). Importantly, less abundant butterflies fit poorly with flowers and the anther mimics. From observations and photographic evidence (***Fig***. *S2B*) the relatively long proboscides of the butterflies visiting the flowers results in the insects probing the flowers between the anther mimics with their heads remaining outside of the flower. As a consequence, butterflies had remarkably low pollen loads and we did not find a single pollen grain from *T. laxifolia* on any of the wings or heads of the 12 butterflies we swabbed for pollen loads. In contrast, *A. fallax* carried 128.9±40.7 SE (n=10) pollen grains on dorsal section of their thorax (*Methods S3*).

The large pollen loads borne by *A. fallax* translated into the substantial number of pollen grains deposited on virgin stigmas in single visit experiments. Virgin stigmas of unmanipulated controls with intact anther mimics received the highest pollen loads compared to control flowers with their anther mimics removed. Few studies have looked at the effects of structural three-dimensional anther mimics in enhancing the morphological fit between flowers and pollinators. The most rigorous studies that do test this, has focused mainly on taxa with vestigial staminodes. For example, Dieringer and Cabrera (2002) found a statistical difference for pollen deposition in *Penstemon digitalis* when comparing control flowers with their staminodes intact, with flowers with their staminodes excised. Similar finding were made by Walker-Larsen and Harder (2001a) for bee pollinated *P. ellipticus* and *P. palmeri*, but not for hummingbird pollinated *P. centranthifolius* and *P. rostriflorus* with their staminodes retained and excised. None of these studies associate pollen collecting behaviour with reproductive success, although there is evidence that the presence of staminodes increase the time spent by pollinators within the flower.

Our videos of the behaviours of bees visiting experimental and control flowers allow us to make a direct link between pollen-collecting behaviour and the amount of contact made with the reproductive organs. Specifically, we found that pollinators displayed a higher proportion of scraping and pulling behaviour on flowers with intact anther mimics (**Fig**. *S5*, *Video S1*).

Reduced pulling behaviours on excised anther mimics is likely the result of a lack of tactile association with the pollen signal. This has fitness implications for two-dimensional pollen imitating markings versus three-dimensional anther mimics with regards to attraction and morphological fit with flowers. Based on our results, pollen imitating markings seem to play an important role in attraction Fig. **2a**), whereas three-dimensional anther mimics may be important in both attraction and in eliciting pollen-collecting behaviour. Ellis and Johnson (2010) showed that ray-florets of the daisy *Gorteria diffusa* with three-dimensional floral signals elicited more mating attempts by male flies compared to plants with two-dimensional floral signals, resulting in more pollen export by the three-dimensional deceptive forms.

In our study, behaviour on three-dimensional anther mimics is associated with a higher proportion of contacts to the reproductive parts of the flower, which is directly linked to pollen deposition and seed set in selection experiments (*see* Newman *et al*., 2015). To our knowledge, this is the first evidence for pollen-collecting behaviour on three-dimensional anther mimics that improves the morphological fit between flower and pollinators (*see* Lunau *et al*. 2017). However, nectar foraging also forces the pollinator to clamber over the anther mimics to access the reward, leading to a higher proportion of contacts to the reproductive organs in unmanipulated controls (**Fig**. 5C). However, pollen collecting behaviour seems to dominate foraging behaviour in the population as unmanipulated flowers received more than two-fold more pollen collecting behaviours compared to nectar foraging behaviours per visit as recorded on video (**Fig**. S5, *Video S1*).

In conclusion, our study makes the link between female reproductive success and the processes underlying the evolution of anther mimicry in *T. laxifolia*. We show that the yellow UV absorbent pollen signal is important in the visual attraction of pollinators to flowers and that the three-dimensional structures not only elicit pollen-collecting behaviours, but such behaviours lead to improved pollen deposition and consequently seed set. Future studies should focus on the generality of pollen-collecting behaviour on two- and three-dimensional pollen imitating structures, and whether inexperienced naïve bees exhibit a higher proportion of pollen collecting behaviour compared to more experienced bees.

## Acknowledgements

We would like to thank the second year Botany class of 2019 of Rhodes University for their enthusiasm for the initial stages of the project as part of their practical assignment. We thank Steve Johnson for comments on an earlier version of this study and Tracey Nowell for bringing us to the attention of the Mosslands population and Barry Hartley for technical assistance. This project was funded by the research council of Rhodes University, South Africa for ELN and CIP.

## Tables and Figures

**Figure 1**. Colour plate of the study system. (a) *Tritonia laxifolia* (Iridaceae) in flower at Kwa-Pikoli, Fish River Pass. (b) *Amegilla fallax* approaches a flower, the white arrow highlights three-dimensional anther mimics on each lower tepal. (c) Bee visitor required to crawl onto and over the anther mimics to contact reproductive parts of the flower. (d) UV images of an unpainted control (left) and a flower with its anther mimics painted with UV reflecting orange paint.

**Figure 2. Figure 2**. Binary preferences of bees based on their first choices (a) between anther mimics painted with yellow UV absorbent paint and orange UV reflecting paint; (b) choices between unpainted controls with naturally yellow anther mimics and anther mimics painted with yellow UV absorbent paint; and (c) choices between unmanipulated controls and flowers with anther mimics excised. Coloured circles represent median model predictions, error bars refer to 90% confidence intervals for model predictions and small coloured points represent balanced binary preferences for each treatment. *ns=P*>0.05, *=*P*<0.05.

**Figure 3**. Colour spectra from different parts of the flower plotted in bee colour space. Colours represent actual colours from respective parts of the flower as perceived by humans. Points with a black outline are measured from anther mimics painted with either orange UV reflecting paint or yellow UV absorbing paint. Spectra in the central grey circle appear achromatic to bees (0.1 hexagon units).

**Figure 4**. Unmanipulated flowers of *Tritonia laxifolia* with anther mimics present, received significantly more pollen deposited on their stigmas following a single visit by a pollinating bee compared to flowers with their anther mimics excised. Coloured circles represent median model predictions, error bars refer to 90% confidence intervals for model predictions and small points represent the number of pollen grains deposited for each single visit replicate. Inset shows full extent of data points. *** *P*<0.001.

**Figure 5**. Pollinator behaviour recorded on unmanipulated controls compared to flowers with anther mimics excised. (a) Proportion of visits with bee scraping anther mimics, (b) Proportion of visits with bees pulling at anther mimics, (c) Proportion of visits by pollen collecting bees (either scraping or pulling behaviour) contacting reproductive organs. Coloured circles represent median model predictions, error bars refer to 90% confidence intervals for model predictions and small points represent binary outcomes. (i.e., presence or absence of behaviour exhibited). **=*P*<0.01*** *P*<0.001.

**Figure 6**. Pollinator behaviour recorded on unmanipulated controls compared to flowers with anther mimics excised. (a) Proportion of visits with bee scraping anther mimics, (b) Proportion of visits with bees pulling at anther mimics, (c) Proportion of visits by pollen collecting bees (either scraping or pulling behaviour) contacting reproductive organs. Coloured circles represent median model predictions, error bars refer to 90% confidence intervals for model predictions and small points represent binary outcomes. (i.e., presence or absence of behaviour exhibited). **=*P*<0.01*** *P*<0.001.

## Supplementary Materials

**Fig. S1.**
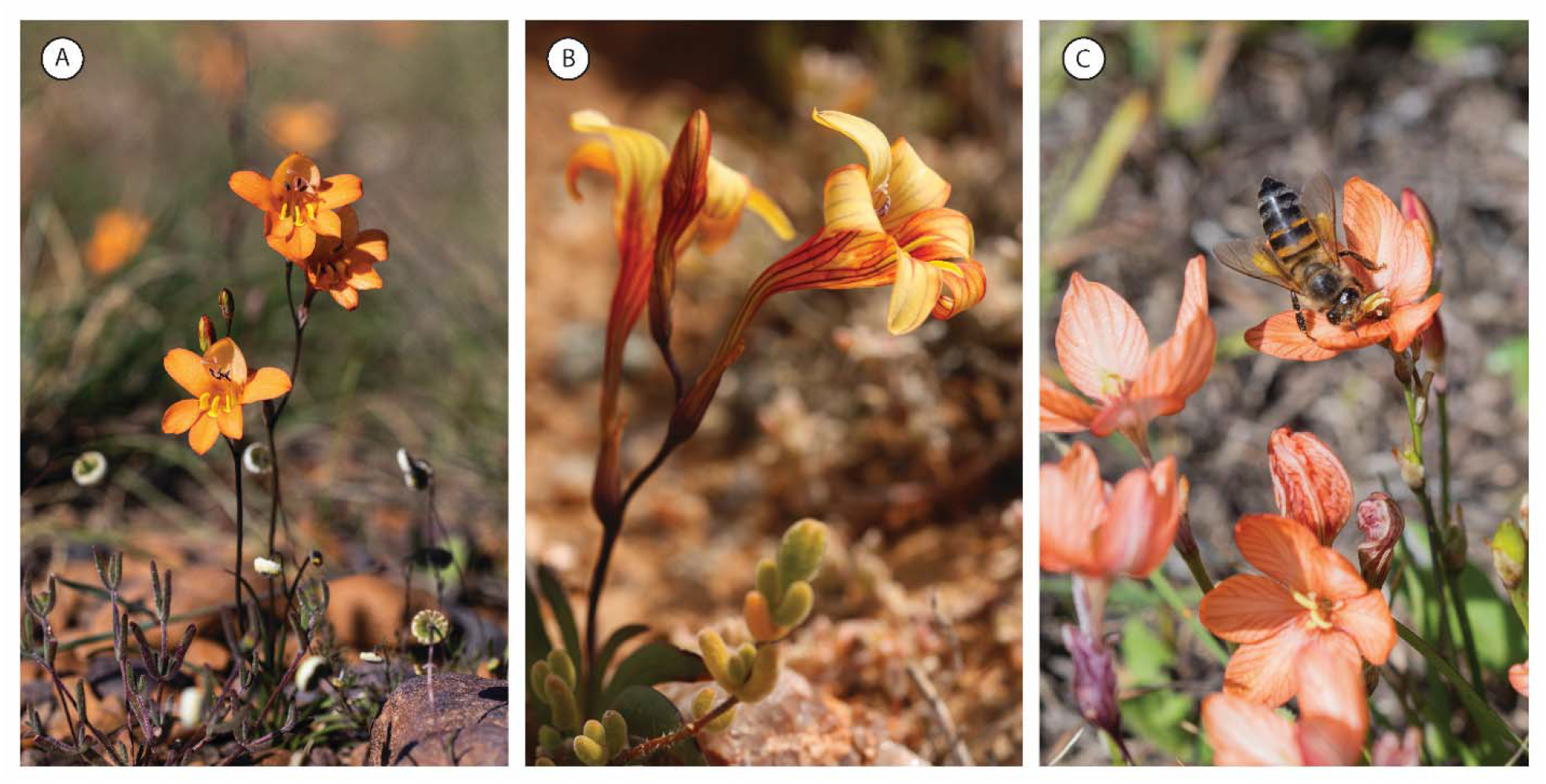
A subset of species from the genus *Tritonia* (Iridaceae) with structural variation in their anther mimics. A) *Tritonia securigera* from Joubertina (Eastern Cape) has “axe-like” three-dimensional anther mimics, B) T. *karooica* from the Roggeveld, (Northern Cape) has slightly raised anther mimics, and C) *T. dubia* from Gqeberha (Eastern Cape) has no anther mimics. Instead, yellow pollen from *T. dubia* anthers attracts honeybees *Apis mellifera scutellata* in search of pollen. All images by Ethan Newman.

**Fig. S2.**
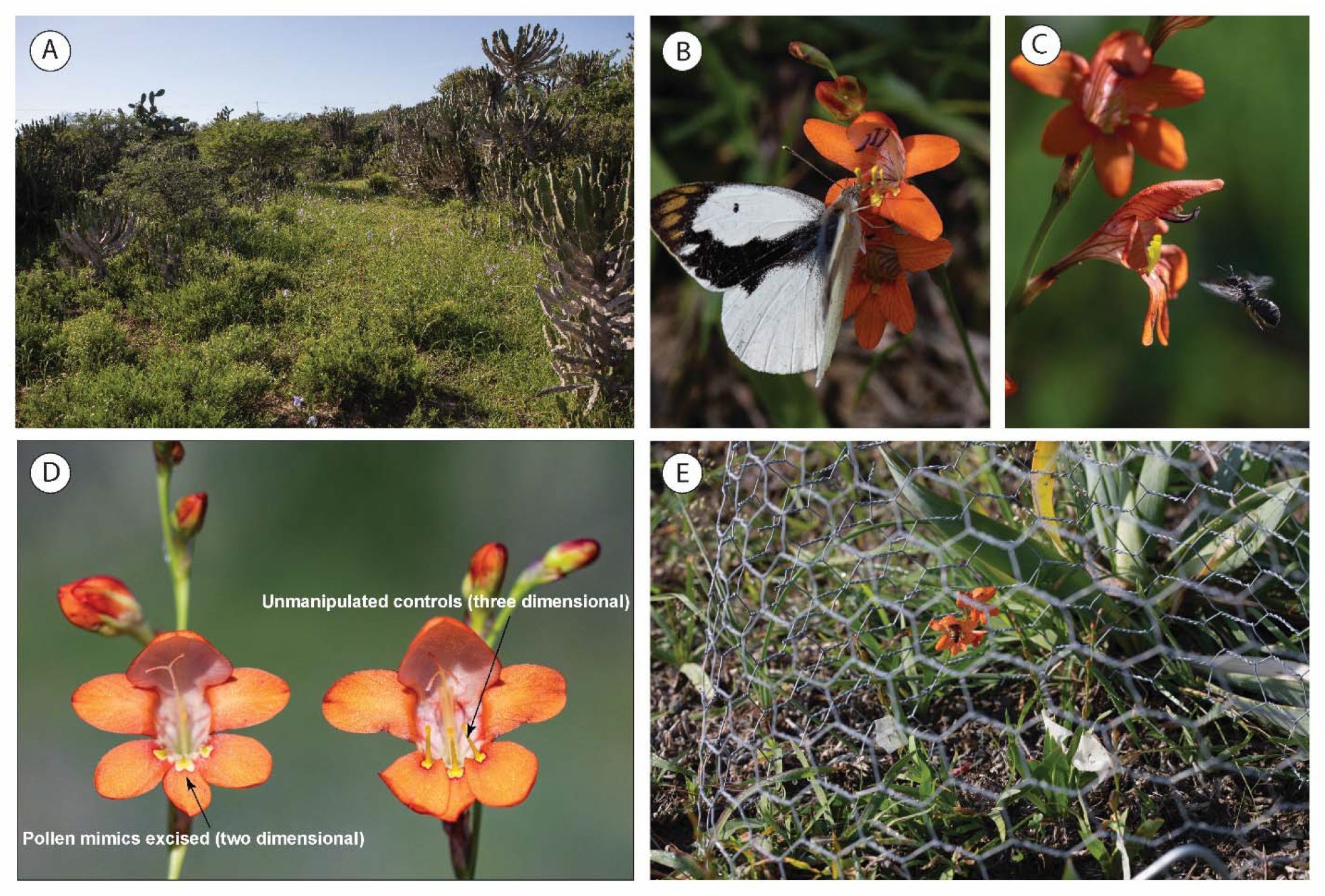
A) Study site where we performed preference and selection experiments near Fish River Pass, Kwa-Pikoli in the Eastern Cape Province of South Africa. B) Gold tip butterfly *Colotis eris* eris thieving nectar from flowers of *Tritonia laxifolia* at the study site. Notice the lack of contact made to the anthers. C*) Allodape* pollen collecting bees approaching a flower of *T. laxifolia* at Makhanda. D) Experimental plants prepared for preference, single visit or selection experiments with the anthers removed in bud and stigmas are receptive, whereby the pollen mimic is excised for one of the treatments, leaving a two-dimensional pollen imitating marking (left), and a flower with the anther mimics kept intact retaining its three-dimensional structure (right). E) Experimental cages placed over rooted plants in the field to exclude butterflies but allowing bees to enter. Notice the bee foraging on the inflorescence inside the cage. All images by Ethan Newman.

**Fig. S3.**
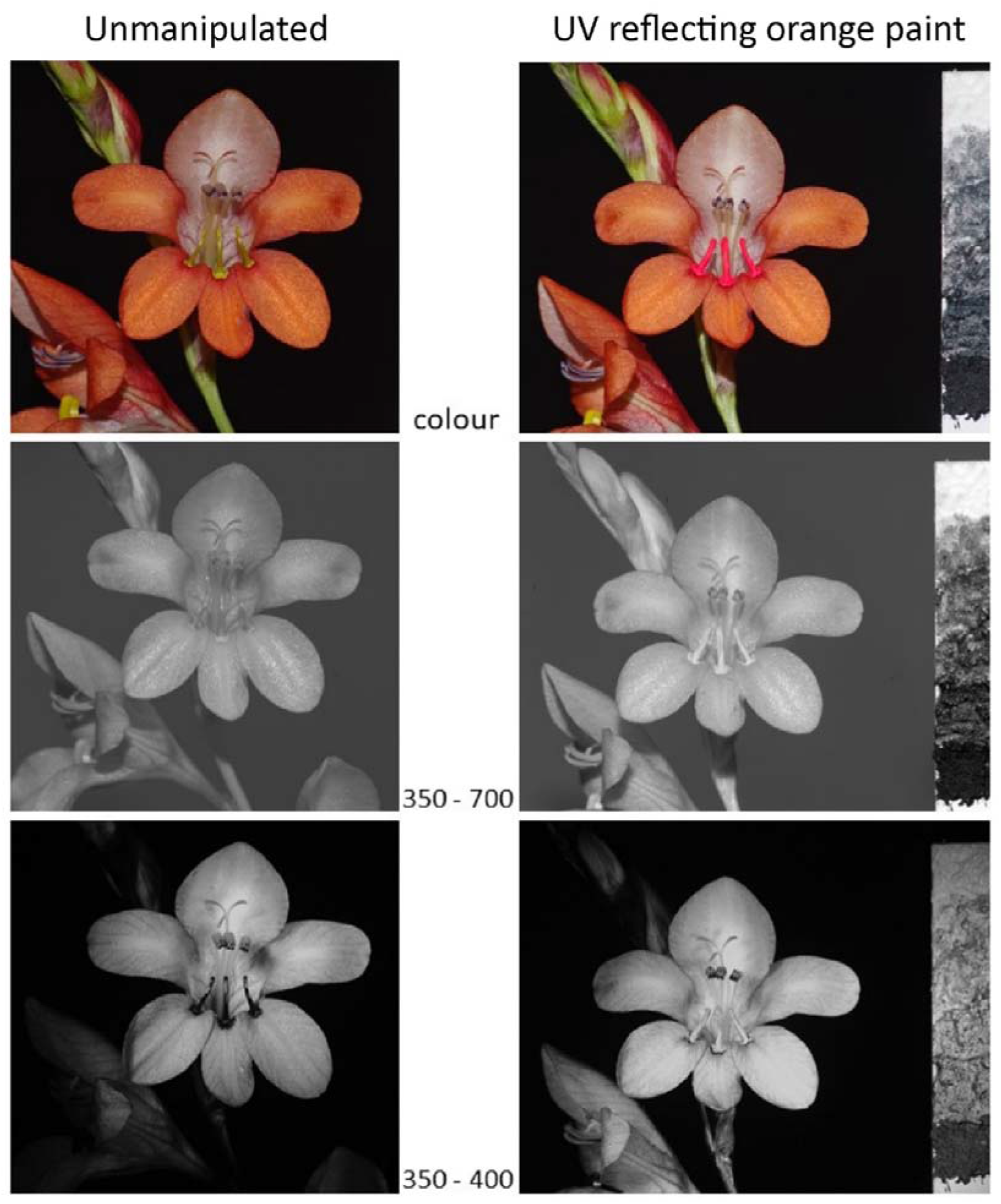
Images on the left represent unpainted and unmanipulated flowers, images on the right have UV reflecting orange paint added to the anther mimics. The first row shows these flowers in colour, the second row between 350 and 700nm, and the last row in the UV range between 350 to 400nm. Notice the UV absorbent properties of the pollen mimic on the unmanipulated flowers on the left versus the UV reflecting paint on the right that appear similar to the tepals. All images by Craig Peter.

**Fig. S4.**
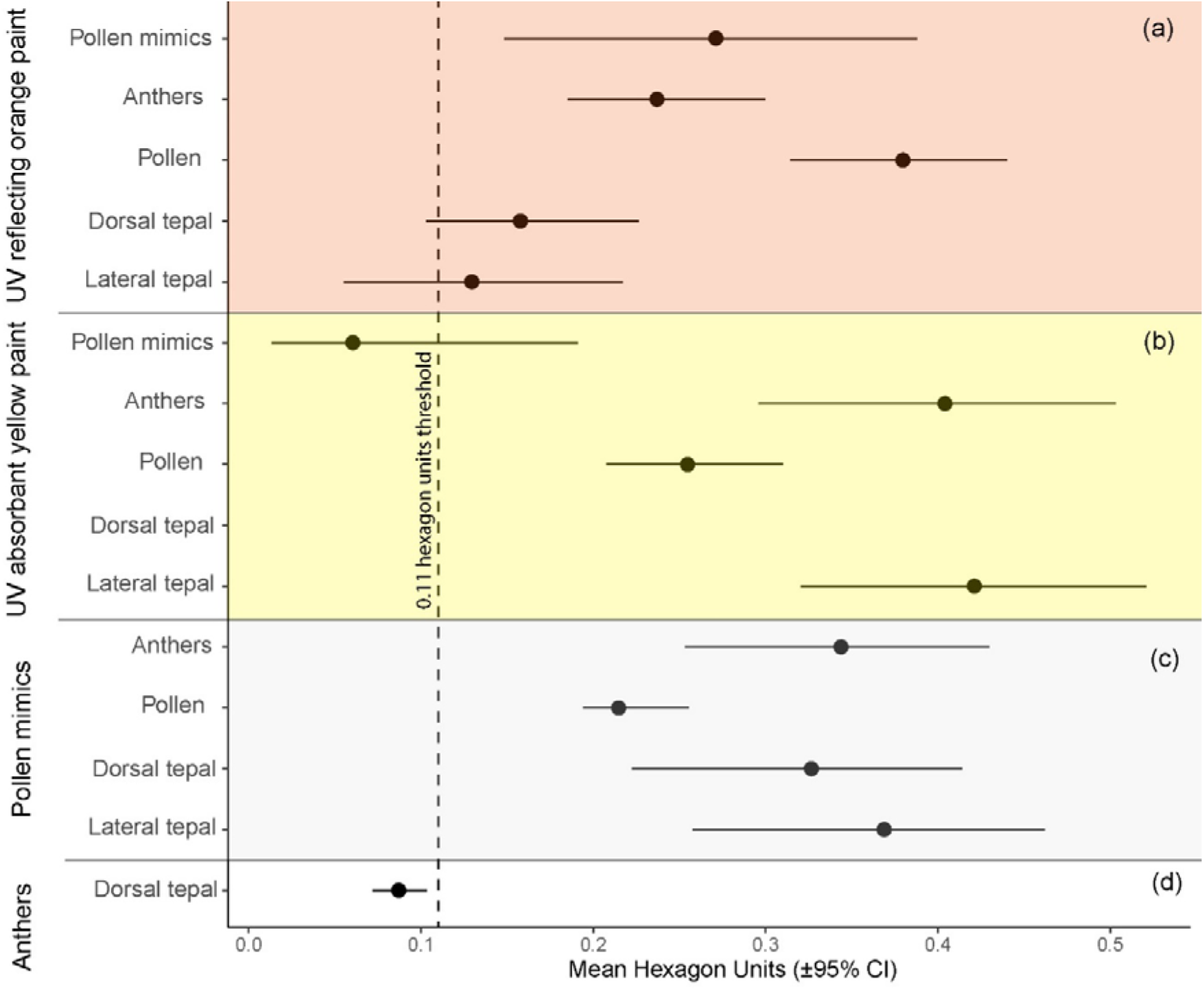
Chromatic contrasts based on Euclidean distances between spectra of different floral traits plotted in bee colour space. (a) UV reflecting orange paint used in colour preferences against floral traits, (b) UV absorbent yellow paint used in colour preferences against floral traits, (c) unmanipulated anther mimics against floral traits. (d) comparison between anthers and dorsal tepals.

**Fig. S5.**
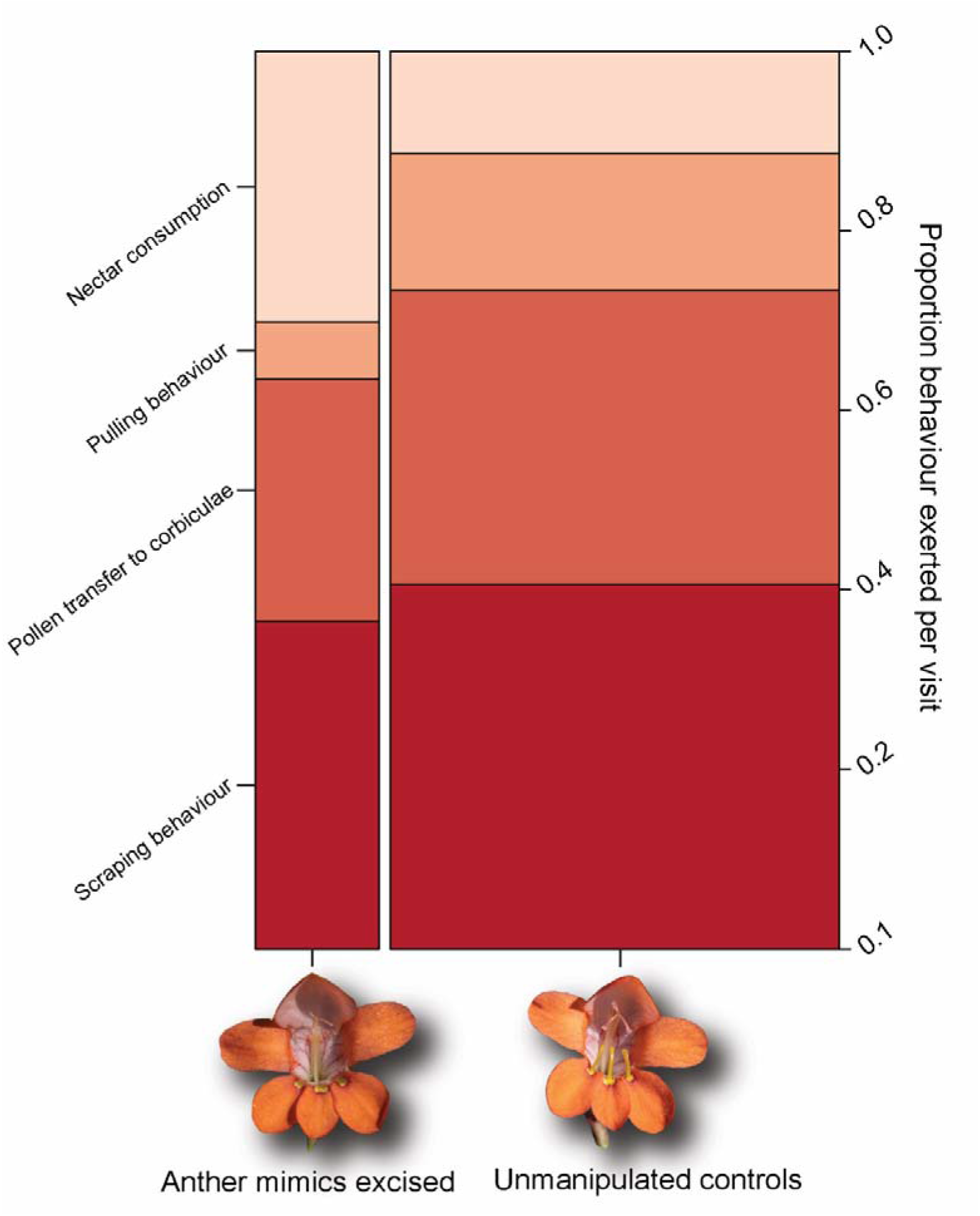
Spine plot illustrating the proportion pollinator behaviour exhibited per single visit by bees. Width of plot represents the number of observations relative to the opposite bar. (i.e., there are more observations of bee behaviour on unmanipulated treatments compared to manipulated treatments).

### Methods S1 UV photography

To assess the qualitative pattern of UV absorbing and UV reflecting parts of the flowers in both unmanipulated flowers and the painted treatments, flowers were photographed in the near UV range (∼350 to 400 nm). This was done using a Pentax K7 subjected to a “full-spectrum” conversion, having the low pass filter covering the sensor removed. A Pentax FA 100mm F3.5 macro lens and a 2-inch Baader Planetarium U-Filter were used, which only transmits wavelengths between 320 and 380 nm. Flowers to be imaged were illuminated with 12V led lights with an emission peak at 365 nm. To judge exposures, a greyscale was constructed using different Magnesium Oxide to Carbon powder ratios and mixed with clear “cold” wood glue to paint the mixture on a white cardboard strip.

### Methods S2 Measurements associated with morphological fit

Floral traits putatively involved in pollination namely tube length, all three calli, the closest distance between the central anther mimic and a stigma branch as well as the measurement between the central lower tepal without the anther mimic and closest stigma branch, were measured from 19 individuals in open receptive female phase flowers. Due to differences in the foraging behaviour of different functional pollinator groups, we measured tube length in two ways. Bees forage for nectar by crawling over the calli and into the flower gullet. Hence, the first measurement was taken in a straight-line distance from the top of the ovary to a notch in the perianth tube which represents the maximum depth that a bee visitor can insert its head required to access the nectar at the bottom. Butterflies on the other hand forage by holding onto the lower tepals and forcing their proboscis through openings between the anther mimics to access the nectar. Hence, flower depth for butterflies was measured as the sum of the first and second measurement, where the second measurement was simply the straight-line distance from the notch in the perianth tube to the furthest distance of the anther mimic with which the head of the butterfly theoretically contacts.

Insect traits potentially important in the pollination process were measured. For Amegilla bees we measured the fully extended proboscis length from the base of the maxilla to the tip of the glossa, a measurement that corresponds with the depth of the perianth tube (tube length measurement 1). Thorax depth was measured as the straight-line distance between the dorsal and ventral portion of the mesothorax. A measurement which corresponds with the distance between the raised nectar guides and the anthers/stigmas. To determine whether butterflies can access nectar from the bottom of the perianth tubes of *Tritonia* flowers, we measured proboscis length from all butterflies as the straight-line distance from the base of the proboscis to the tip, by extending the knee bend. A measurement that corresponds with the depth of the perianth tube (The sum of tube length measurements 1 and 2). All measurements were taken using a set of digital callipers calibrated to 0.1 millimeter.

### Methods S3 Determining the number of pollen grains on pollinators

During pollinator preference experiments, one of the observers (ELN) walked through the population at Fish River Pass and caught all insects visiting *T. laxifolia* flowers. On the 19^th^ of May 2019 we captured all visitors actively foraging on *T. laxifolia* by insect net and killed them by freezing them on site. Bees were placed in a tapering 150 ml tube with their headfirst, to restrict their movements as much as possible to prevent pollen from falling off and moving to different parts of their bodies. Butterflies were killed by gently squeezing their thorax following capture, inserting them in resting position in a wax paper envelope (Bio Quip Products. California, USA) before freezing.

To assess the degree of morphological fit of flowers with different functional pollinator groups, we counted the number of *T. laxifolia* pollen grains exported on the bodies of captured insects. *Tritonia laxifolia* pollen was counted from 12 Anthophorid bees (*Amegilla sp*.), 11 Gold tip butterflies (*Colotis eris eris*). For each individual insect, fuchsin gel was gently dabbed onto the area of pollen deposition. For bees, fuchsin gel was dabbed onto the top of the bees’ head, dorsal section of the thorax and forewings and for butterflies, fuchsin gel was dabbed onto the proboscis, top of head and first half of the dorsal section of the thorax. Using reference slides, *T. laxifolia* pollen was discerned from pollen of all plant species in the community visited by the respective visitors to *T. laxifolia*.

**Video/Movie S1** Video clip illustrating pollen-collecting behaviour exerted on anther mimics by bees

## Notes

### Competing Interest Statement

The authors have declared no competing interest.

